# Multiple ER-to-nucleus stress signaling pathways become active during *Plantago asiatica mosaic virus* and *Turnip mosaic virus* infection in *Arabidopsis thaliana*

**DOI:** 10.1101/786137

**Authors:** Mathieu Gayral, Omar Arias Gaguancela, Evelyn Vasquez, Venura Herath, Mingxiong Pang, Francisco Javier Florez, Martin B Dickman, Jeanmarie Verchot

## Abstract

Endoplasmic reticulum (ER) stress due to biotic or abiotic stress activates the unfolded protein response (UPR) to restore ER homeostasis. The UPR relies on multiple ER-to-nucleus signaling factors which mainly induce the expression of cytoprotective ER-chaperones. The inositol requiring enzyme (IRE1) along with its splicing target, bZIP60, restrict potyvirus, and potexvirus accumulation. Until now, the involvement of the alternative UPR pathways and the role of UPR to limit virus accumulation have remained elusive. Here, we used the *Plantago asiatica mosaic virus* (PlAMV) and the *Turnip mosaic virus* (TuMV) to demonstrate that the potexvirus triple gene block 3 (TGB3) protein and the potyvirus 6K2 protein activate the bZIP17, bZIP28, bZIP60, BAG7, NAC089 and NAC103 signaling in *Arabidopsis thaliana*. Using the corresponding knock-out mutant lines, we demonstrated that these factors differentially restrict local and systemic virus accumulation. We show that bZIP17, bZIP60, BAG7, and NAC089 are factors in PlAMV infection, whereas bZIP28 and bZIP60 are factors in TuMV infection. TGB3 and 6K2 transient expression in leave reveal that these alternative pathways induce BiPs expression. Finally, using dithiothreitol (DTT) and tauroursodeoxycholic acid (TUDCA) treatment, we demonstrated that the protein folding capacity significantly influences PlAMV accumulation. Together, these results indicate that multiple ER-to-nucleus signaling pathways are activated during virus infection and restrict virus accumulation through increasing protein folding capacity.

**Significance statement:** The IRE1/bZIP60 pathway of unfolded protein response (UPR) is activated by potyviruses and potexviruses, limiting their infection, but the role of alternative UPR pathways is unknown. This study reveals the activation of multiple ER-to-nucleus signaling pathways by the *Plantago asiatica mosaic virus* and the *Turnip mosaic virus.* We identify additional signaling pathways serve to restrict virus accumulation through increased protein folding capacity.

## Introduction

The endoplasmic reticulum (ER) and Golgi network house the cellular machinery for protein synthesis, maturation, and delivery to the intended subcellular compartments. Environmental challenges such as heat, drought, and pathogen attack can drastically alter the protein maturation process and cause a bottleneck of malformed proteins to accumulate in the ER. The unfolded protein response (UPR) describes alternative ER-to-nucleus signaling pathways leading to increased nuclear gene expression (Bao *et al.*, 2018, Nawkar *et al.*, 2018).

Transmembrane sensors embedded in the ER respond to the accumulation of aberrant proteins to increase the synthesis of chaperones and the endoplasmic-reticulum-associated protein degradation machinery to refold or decay proteins to alleviate any bottleneck of protein maturation in the ER and recover ER homeostasis (Angelos *et al.*, 2017). When ER stress is chronic, and cells cannot return to homeostasis, there is heightened autophagy and cell death (Srivastava *et al.*, 2018).

One pathway in plants involves the inositol-requiring enzyme (IRE1) and the basic leucine zipper 60 (bZIP60) transcription factor. *Arabidopsis thaliana* has two copies of *IRE1* known as *IRE1a* and *IRE1b* that catalyze the splicing of *bZIP60* mRNAs (Deng *et al.*, 2013). In the nucleus, the truncated bZIP60 transcription factor upregulates the expression of several ER-resident chaperones including *protein disulfide isomerase (PDI), calnexin (CNX), calreticulin (CRT) and the HSP70-like binding immunoglobulin protein (BiP)* (Iwata *et al.*, 2008). A separate branch of the IRE1-led pathway targets mRNAs of secreted proteins in a process called Regulated IRE1-Dependent Decay of mRNAs (RIDD). The RIDD activity of IRE1b is linked to the activation of autophagy in response to ER stress (Liu *et al.*, 2012, Bao *et al.*, 2018).

The second major pathway is mediated by bZIP17 and bZIP28. Under normal conditions, bZIP17 and bZIP28 reside in the ER membrane (Angelos *et al.*, 2017, Kim *et al.*, 2018). The Bcl-2–associated athanogene 7 (BAG7) is an ER-localized co-chaperone that under regular conditions associates with bZIP28 and BiP in the ER (Williams *et al.*, 2010, Li *et al.*, 2017) (Srivastava *et al.*, 2013). The BAG7 and bZIP28 can be activated in cells following heat stress or chemical stress, while bZIP17 is reported to be activated by salt stress. Upon activation, the BAG7/bZIP28/BiP complex dissociates, and the bZIP28 and BAG7 are available to move between the ER and the nucleus(Williams *et al.*, 2010, Li *et al.*, 2017). The SITE-1 Protease (S1P) and S2P remove the transmembrane domains releasing the bZIP17 and bZIP28 transcription factors to enter the nucleus (Liu *et al.*, 2007b, Liu *et al.*, 2007a, Iwata *et al.*, 2017) and activate expression of BiPs and other chaperones, similar to bZIP60 (Nawkar *et al.*, 2018). The transmembrane domain of BAG7 is also proteolytically removed, and the truncated BAG7 translocates to the nucleus where it principally interacts with WRKY28, independent of bZIP28. The BAG7/WRKY28 induce transcription of stress-responsive chaperones, including BAG7 itself, which protects the cell and ensures its survival (Li *et al.*, 2017).

Generally, bZIP17, bZIP28, and bZIP60 can form homo- and heterodimers (Deppmann *et al.*, 2004) that further increase the possible regulatory combinations. The bZIP factors recognize G-box core promoter sequences (CACGTG). The flanking sequences near each G-box determines recognition specificity by each factor (Ezer *et al.*, 2017). We know very little about which stretches of sequences containing G-boxes are differential targets for homo-and heterodimers of these bZIP factors and cannot yet precisely predict the transcription pattern of genes by examining promoter sequences (Ezer *et al.*, 2017). Nevertheless, experiments have demonstrated that the bZIP60 and bZIP28 recognize the promoters of at least two NAC (NAM/ATAF/CUC) transcription factors (Nawkar *et al.*, 2018). *NAC103* is activated by bZIP60 through the UPRE-III *cis*-elements and upregulates the expression of genes that are categorically described as “pro-survival” factors (Sun *et al.*, 2013). NAC089 is localized in the ER membrane and is regulated by bZIP28 and bZIP60 heterodimers through the UPRE and ERSE-I *cis*-elements (Yang *et al.*, 2014). *The* silencing of *NbbZIP28* or *NbNAC089* in *Nicotiana benthamiana* increases plant susceptibility to the *Tobacco mosaic virus* (TMV) and *Cucumber mosaic virus* (CMV) (Shen *et al.*, 2017, Li *et al.*, 2018). During ER stress, NAC089 promotes transcriptional activation of *BAG6* (Yang *et al.*, 2014). BAG6 is a co-chaperone with a calmodulin-binding domain and can activate autophagy required for basal immunity against the necrotrophic fungus *Botrytis cinerea* (Li *et al.*, 2016).

Upon entry into the host cells, *Plantago asiatica mosaic virus* (PlAMV) (genus: *Potexvirus*) and *Turnip mosaic virus* (TuMV) (genus: *Potyvirus*), induce rearrangements of organellar membranes, including the ER (Verchot, 2016). Viruses principally use the ER as a replication site and guide for cell-to-cell movement, forming vesicles or spherules surrounding viral replication complexes (Tilsner *et al.*, 2013, Cabanillas *et al.*, 2018). These vesicles/spherules concentrate viral proteins necessary for replication and protection against cellular defenses. The ER is essential for potyviruses and potexviruses as a guide for cell-to-cell movement through plasmodesmata (Verchot-Lubicz *et al.*, 2010, Grangeon *et al.*, 2013). The movement from cell-to-cell permits the virus to spread inside the leaves, called local infection. Then virus loads into sieve elements and spreads throughout the plant via the phloem, called systemic infection. The potyvirus membrane-binding protein 6K2 and potexvirus triple gene block 3 (TGB3) are essential in these mechanisms. Several studies reported that potexviruses and potyvirus infection induce ER stress and the related UPR signaling (Ye *et al.*, 2011, Ye *et al.*, 2013, Verchot, 2014, Zhang *et al.*, 2015, Gaguancela *et al.*, 2016, Verchot, 2016). Each viral protein of TuMV and *Potato virus Y* (PVY)(genus: *Potyvirus*) as well as PlAMV and *Potato virus X* (PVX) (genus: *Potexvirus*) protein, was transiently expressed in *N. benthamiana* or Arabidopsis leaves using agro-delivery of binary constructs and it was demonstrated that the potyvirus 6K2 and potexvirus TGB3 proteins function as viral inducer of UPR via the IRE1/bZIP60 pathway (Zhang *et al.*, 2015, Gaguancela *et al.*, 2016). We also used infectious clones of *PlAMV and TuMV* which have GFP introduced into the viral genomes to inoculate plants and investigate how these viruses interact with the UPR machinery (Ye *et al.*, 2011, Ye *et al.*, 2013, Gaguancela *et al.*, 2016). GFP expression from the viral genome can be used to visualize and quantify virus accumulation in knockout (KO) mutant plants that have defects in the UPR machinery and wild-type Col-0 plants. This powerful technology has enabled us to demonstrate that PlAMV-GFP and TuMV-GFP accumulate to higher levels in local and systemic leaves in *ire1a/ire1b* and *bzip60* mutants than wild-type Col-0 plants (Zhang *et al.*, 2015, Gaguancela *et al.*, 2016). Moreover, we reported that TGB3 preferentially engages IRE1a, whereas 6K2 preferentially engages IRE1b in Arabidopsis plants. In this study, we investigate whether bZIP17 and bZIP28 are engaged in UPR responses to infection by PlAMV-GFP and TuMV-GFP in Arabidopsis plants. Our results reveal that multiple ER-to-nucleus stress signaling pathways respond to TGB3 and 6K2 proteins. These alternative UPR pathways converge to mainly restrict PlAMV-GFP and TuMV-GFP accumulation in plants through enhancing the protein folding capacity of the ER.

## Results

### Potexvirus TGB3 and potyvirus 6K2 induce expression of *bZIP60, bZIP28,* and *bZIP17*

Increased accumulation of *bZIP60, bZIP28, or bZIP17* transcripts is typically reported as evidence of UPR activation (Iwata and Koizumi, 2012, Fanata *et al.*, 2013). Since research has well established that the potexvirus TGB3 and potyvirus 6K2 proteins activate bZIP60-led upregulation of UPR related genes, we investigated whether bZIP28 and bZIP17 led branch pathways respond to delivery of these viral proteins. We established that quantitative RT-PCR (qRT-PCR) could be used to detect changes in transcript levels of bZIP60 and other UPR related chaperones at 2- and 5-days post infiltration (dpi) following agro-delivery of binary vectors expressing the PlAMV TGB3, PVX TGB3, PVY 6K2, or TuMV 6K2 genes (Ye *et al.*, 2011, Ye *et al.*, 2013, Zhang *et al.*, 2015). In this study, binary constructs expressing PlAMV TGB3, PVX TGB3, PVY 6K2, and TuMV 6K2 were delivered by agro-infiltration to individual Col-0 leaves (n=3 plants). The treatment of mock controls was with *A. tumefaciens* cultures that did not contain the constructs expressing the viral genes. Multiple infiltrated leaves were harvested from each plant and pooled for RNA extraction at 2 and 5 dpi. The qRT-PCRs were performed with the three experimental replicates to obtain the average level of RNA accumulation relative to the mock-treated control within each experiment. Experiments were repeated multiple times to confirm the reproducibility of individual experiments. At two dpi, *bZIP17, bZIP28*, and *bZIP60* expression levels were between 2- and 11-fold higher than the control, following expression of each viral *TGB3* or *6K2* gene (Figure 1a; P<0.05). At 5 dpi, *bZIP60* expression levels remained elevated in all samples (P<0.05) while *bZIP17* transcripts declined to control levels for most treatments except in PVX TGB3 treated leaves (Figure 1b; P<0.05). The level of *bZIP28* transcripts also dropped but remained above the control levels in PlAMV TGB3, and PVY 6K2 treated leaves (Figure 1b; P<0.05).

**Figure 1.**
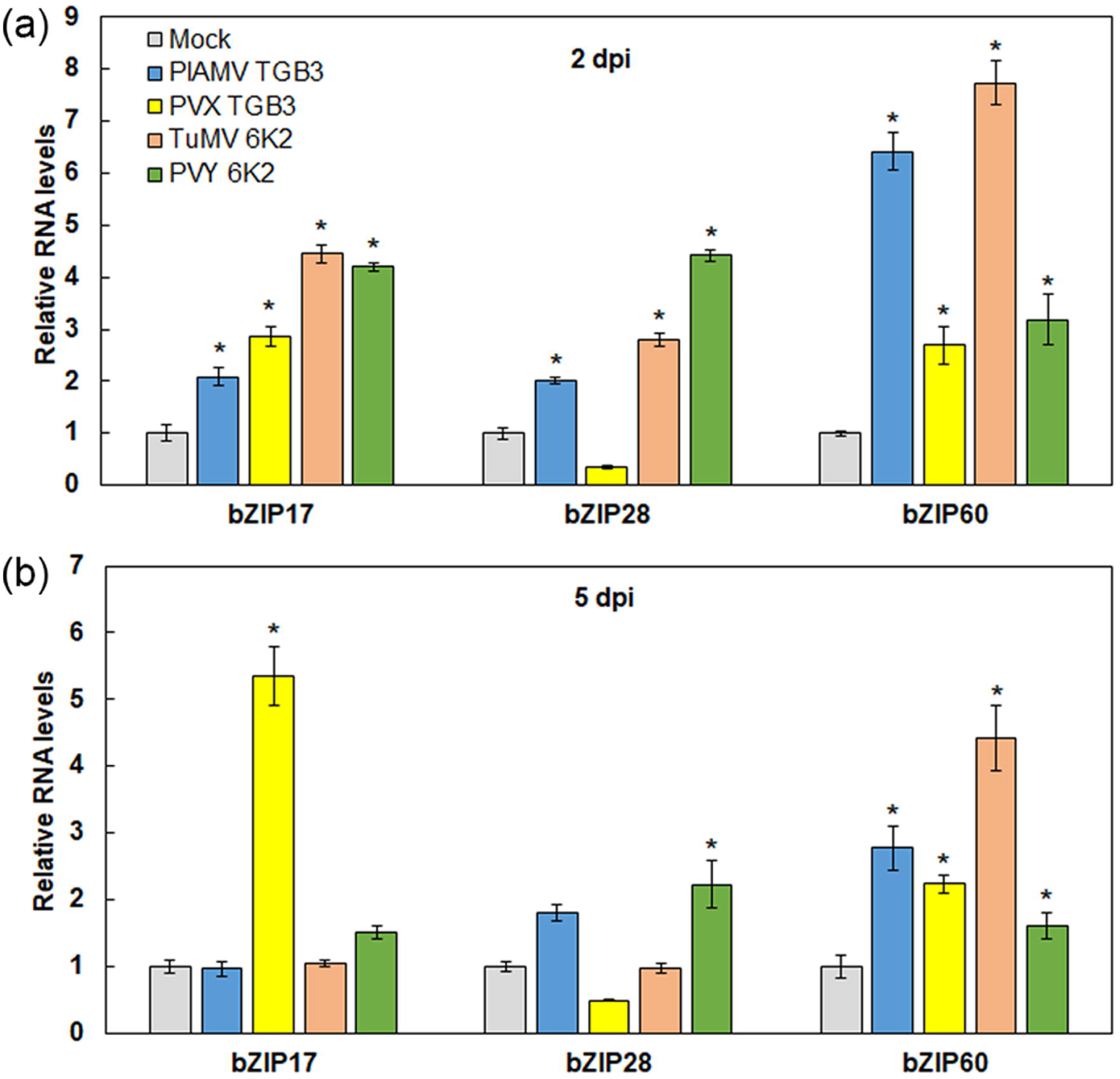
TGB3 and 6K2 induces transcription of *bZIP17 bZIP28* and *bZIP60* genes in *Arabidopsis* Col-0 leaves. qRT-PCR results are providing relative RNA levels of *bZIP17, bZIP28,* and *bZIP60* following treatments with each viral factor at 2 (a) and 5 dpi (b). Error bars represent the standard deviation (SD); asterisks indicate significant differences to the mock control: Student’s *t*-test; *P*<0.05; n=3.

### bZIP60 and bZIP17 significantly reduce PlAMV-GFP accumulation

We conducted a series of experiments to learn whether bZIP28 and bZIP17 play a role in local infection by promoting or restricting virus accumulation in the inoculated leaves. PlAMV-GFP infectious clones were used to inoculate *bzip60, bzip17, bzip28, bzip60/bzip17* and *bzip60/bzip28* KO lines, as well as wild-type Col-0 plants. Immunoblot detection of the viral coat protein (CP), as well as GFP fluorescence were used to detect virus infection in the inoculated leaves. We examined leaves every day and determined that GFP fluorescence appears at 4 dpi. We identified 5 dpi as the optimum time to isolate leaves for analysis (Figure S1). Immunoblots revealed higher levels of CP in *bzip60, bzip17, bzip60/bzip17, and bzip60/bzip28* leaves than in *bzip28* or Col-0 leaves (Figure 2a). Image J was used to quantify GFP fluorescence value (FV) in individual leaves (n=6). Based on the average FVs, PlAMV-GFP accumulation was significantly higher in the *bzip60, bzip17, bzip60/17* and *bzip28/bzip60* leaves (between 3.1- and 9.4-fold; P<0.05) than in the Col-0 leaves (Figure 2b). The FV on *bzip28* leaves were not different from Col-0 (Figure 2b). These data suggest that bZIP60 and bZIP17 significantly contribute to restricting PlAMV-GFP local infection.

**Figure 2.**
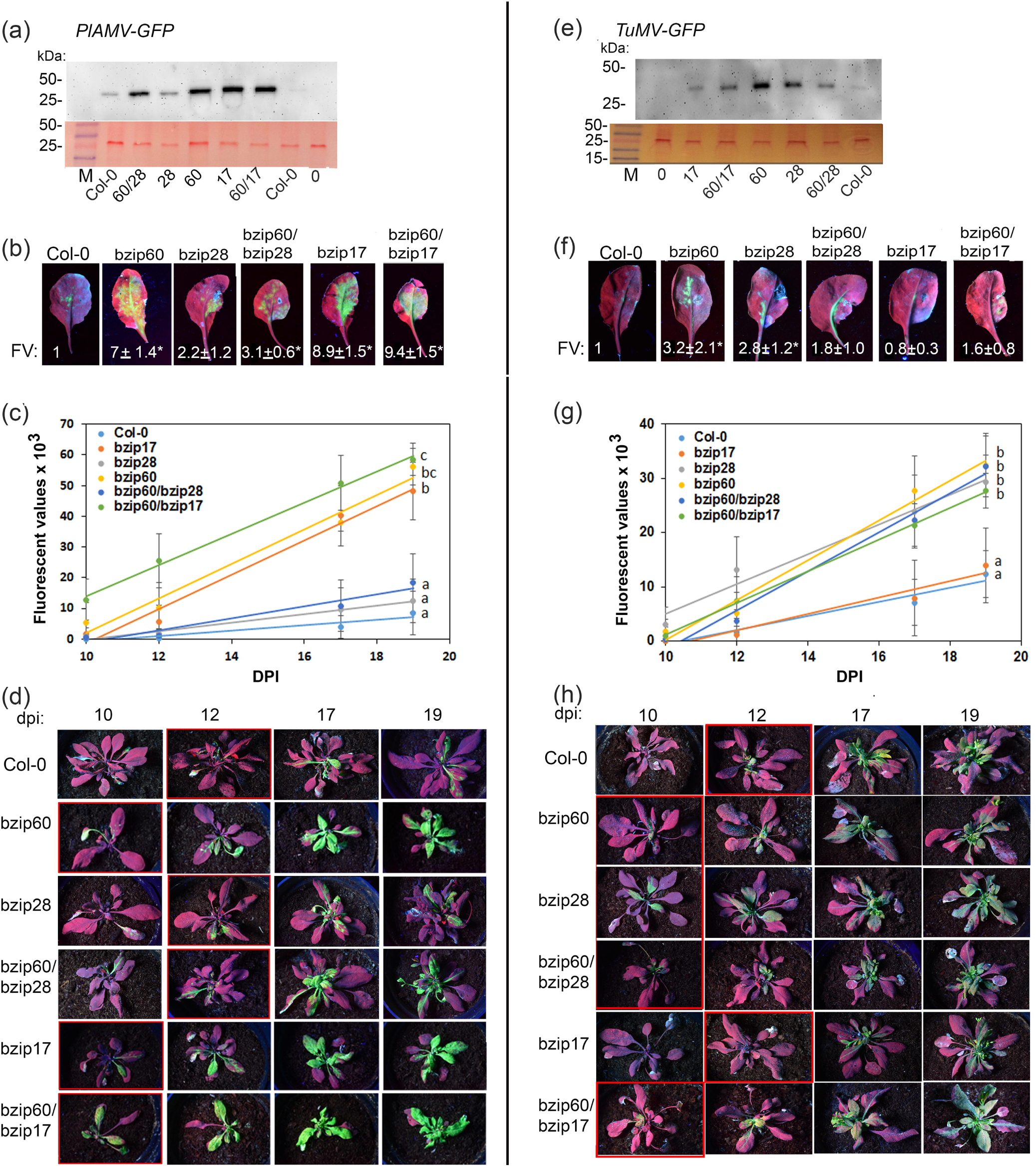
Enhanced local and systemic infection of PlAMV-GFP and TuMV-GFP in *bzip60, bzip17, bzip28*, and double mutant plants compared to Col-0 plants. Immunoblots are detecting PlAMV (a) and TuMV (e) CPs in the inoculated leaves at 5 dpi. Ponceau S-stained membranes below the blots show equal protein loading. The labels identify each *bzip* mutant host below each lane. M= molecular weight marker; “0” is mock. Representative images of PlAMV-GFP (b) and TuMV-GFP (f) inoculated leaves at 5 and 8 dpi respectively. The average fluorescence value (FV) and SD is reported below each image, and the asterisks indicate statistical differences: Student’s *t*-test; P<0.05; n=6. Scatter plots show the average FVs, SDs, and trend lines for mutant and Col-0 plants inoculated with PlAMV-GFP (c) and TuMV-GFP (g). Letters on the right of each line represents the statistical relatedness of the FVs for each treatment and at each time point as determined by ANOVA; *P*<0.05; n=6. Representative images of plants infected with PlAMV-GFP (d) and TuMV-GFP (h) at 10, 12, 17, and 19 dpi. Red boxes surround the plants at 10 or 12 dpi showing the first signs of systemic fluorescence, which in some cases was a small area of a single leaf and in other cases was a broad area.

To learn if these bZIP factors play a role in promoting or restricting systemic infection, we first conducted a time course to study GFP systemic accumulation until 19 days. GFP fluorescence in Col-0 plants unloaded from the midrib and primary branching veins into the first upper leaves at 12 dpi, and the fluorescence spread throughout several upper leaves 19 days (Figure S1). While there was a continued spread of GFP beyond 19 dpi, the time between 10 and 19 dpi provided a linear range for GFP FVs that allowed comparisons between wild-type and KO lines. By including 10 dpi as a time point, we were able to determine if the systemic infection was earlier or delayed in the KO lines compared to the Col-0 plants. In these and subsequent experiments, Image J was used to quantify systemic GFP fluorescence images taken at 10, 12, 17, and 19 dpi (n=6). The FVs were plotted and statistically analyzed (Figure 2c; P<0.05). The systemic FVs due to PlAMV-GFP increased at a high rate in *bzip60, bzip17*, and *bzip60/bzip17* plants and were statistically different from the average FVs calculated in Col-0 plants (Figure 2c; P<0.05). It is interesting to note that PlAMV-GFP accumulation is significantly higher in *bzip60/bzip17* than in *bzip17* plants. However, there was no significant difference between systemic FVs in *bzip60* and *bzip60/bzip17* plants, suggesting that the bZIP60 and bZIP17 act independently. The average FVs in Col-0, *bzip28,* and *bipz60/bzip28* plants were not significantly different from each other (Figure 2c; P<0.05). We first observed GFP at 10 dpi in the upper leaves of *bzip60, bzip17,* and *bzip60/zip17* plants but at 12 dpi in Col-0, *bzip28*, and *bzip60/28* plants (Figure 2d). These combined results indicate that bZIP60 and bZIP17 contribute to restricting the local and systemic infection of PlAMV-GFP, whereas bZIP28 does not limit PlAMV-GFP infection.

### bZIP60 and bZIP28 significantly reduce TuMV-GFP accumulation

We conducted similar experiments to learn whether bZIP28 and bZIP17, alongside bZIP60, promote or restrict TuMV-GFP local and systemic infection. We examined GFP fluorescence each day to identify the optimum duration to measure GFP and coat protein levels that is nearest to the time of gene induction, as direct indicators of virus accumulation in wild type and mutant plants. The onset of TuMV-GFP infection in the inoculated leaves is slower than PlAMV-GFP. Although we first observed GFP at 6 dpi, we identified 8 dpi was preferable for immunoblot analysis of CP levels and fluorescence quantification (Figure S1). Virus CP accumulation was higher in *bzip60, bzip28, bzip60/bzip17*, and *bzip60/bzip28* infected leaves than in Col-0 infected leaves (Figure 2e). The average FVs in the TuMV-GFP inoculated leaves of *bzip60*, and *bzip28* plants were significantly higher than in Col-0 plants (Figure 2f; P<0.05, n=6), whereas the average FVs in *bzip17* leaves were not different from the average FVs in Col-0 leaves. These data suggest that *bZIP60* and *bZIP28*, but not *bZIP17*, contribute to establishing local infection.

To learn if these bZIP factors play a role in promoting or restricting TuMV-GFP systemic infection, we conducted time-course experiments to study GFP systemic accumulation over 19 days. GFP fluorescence in Col-0 plants appeared to be unloading into the first upper leaves at 12 dpi, and the fluorescence continued to spread through 19 dpi, similar to PlAMV-GFP (Figure S1). We measure GFP fluorescence at 10, 12, 17, and 19 dpi using the Image J software, and the average FVs were statistically compared (n=6). The average GFP FVs were significantly higher in *bzip60, bzip28, bzip60/bzip28*, and *bzip60/bzip17* plants than in *bzip17* or Col-0 plants (Figure 2g; P<0.05). It is interesting to note that *bzip60, bzip28, bzip60/bzip28* are not significantly different, suggesting that effects of the mutations in bZIP60 and bZIP28 are not additive. At 10 dpi, GFP appeared systemically in *bzip60, bzip28, bzip60/bzip28* and *bzip60/bzip17* plants, but appeared at 12 dpi in the upper leaves of *bzip17* and Col-0 plants (Figure 2h). Finally, these results reveal that bZIP60 and bZIP28 restricts TuMV-GFP local and systemic infection.

### BAG7 reduces the local accumulation of PlAMV-GFP but not TuMV-GFP

BAG7 is a recently discovered hallmark of UPR during heat stress and is involved in two separate activities. First, BAG7 is engaged in the retention of bZIP28 in the ER. Second, BAG7 translocates to the nucleus where it participates with WRKY29 to upregulate genes involved in heat stress resistance (Li *et al.*, 2017). We hypothesized that if bZIP28 plays a role in restricting TuMV-GFP but not PlAMV-GFP, then BAG7 may also be engaged in regulating virus accumulation. First, Col-0 and *bag7* plants were inoculated with PlAMV-GFP and at 5 dpi, the average FV was 6.9-fold higher in the *bag7* than in the Col-0 inoculated leaves (Figure 3a; P<0.05; n=6). Immunoblots also showed that the CP levels were relatively higher in *bag7* compared to Col-0 leaves (Figure 3a). These data indicate that BAG7 is engaged in restricting PlAMV-GFP local infection. As in previous experiments, we recorded and statistically analyzed FVs in systemic leaves between 10 and 19 dpi (Figure 3b; n=12). In these experiments, there was no difference in FVs in the upper leaves of Col-0 and *bag7* plants (Figure 3b and c; P<0.05). These results demonstrated that BAG7 is essential for restricting PlAMV-GFP local infection in the inoculated leaves but does not limit systemic infection.

**Figure 3.**
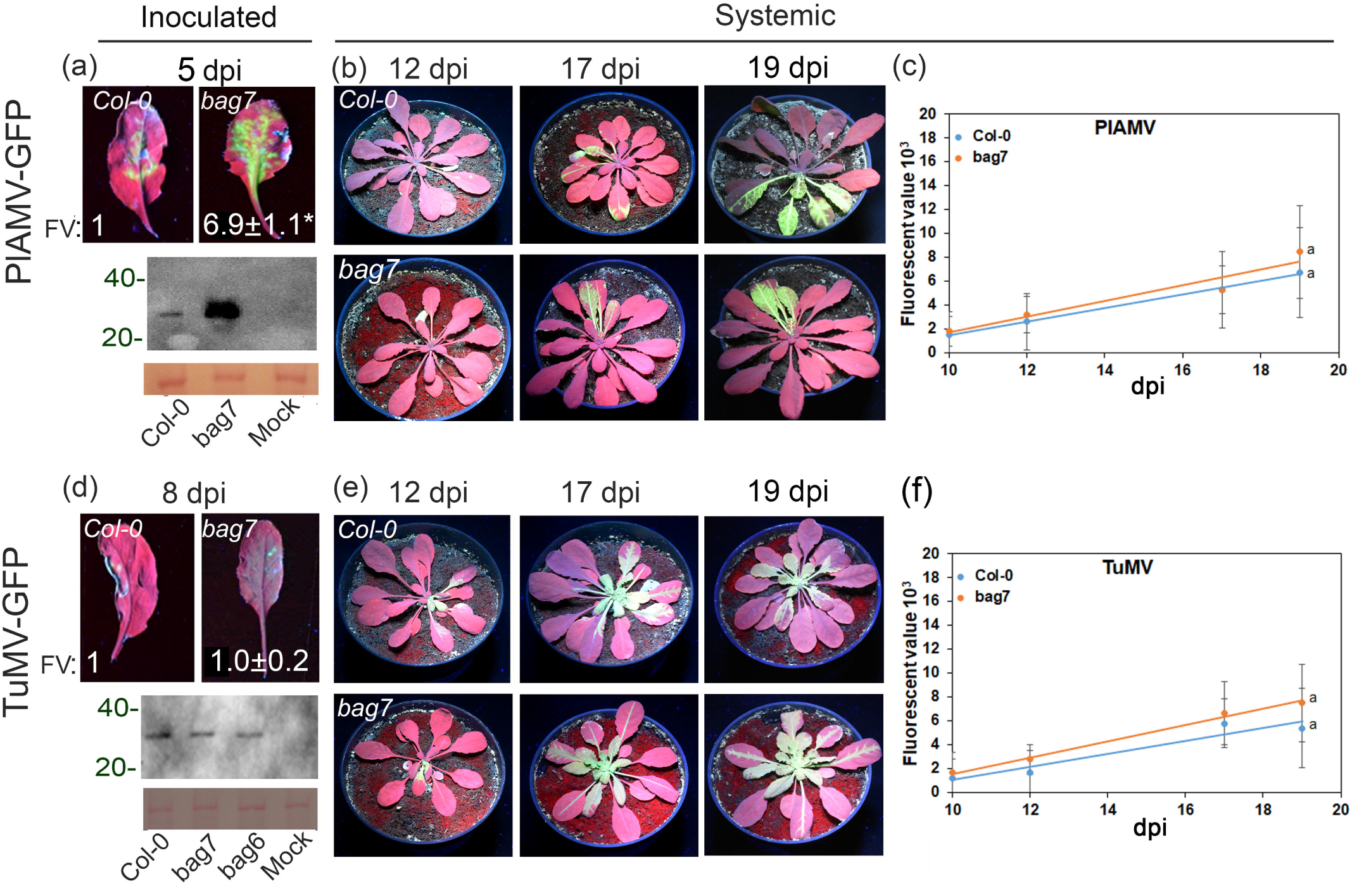
Local and systemic infection of PlAMV-GFP and TuMV-GFP in *bag7* and Col-0 plants. Representative images of PlAMV-GFP (a) and TuMV-GFP (d) inoculated leaves at 5 and 8 dpi, respectively. The average FVs, the SDs, and statistical relatedness are reported at the bottom of each panel. Asterisks indicate statistical differences: Student’s *t*.test; P<0.05; n=6. Immunoblot analysis detecting PlAMV (a) and TuMV (d) CPs in the inoculated leaves, respectively. Ponceau S-stained membrane below the immunoblot shows total equal protein loading. Representative images of plants infected with PlAMV-GFP (b) and TuMV-GFP (e) at 12, 17and 19 dpi. Scatter plot values and trend lines represent the average FV and SDs for each mutant and Col-0 line inoculated with PlAMV-GFP (c) and TuMV-GFP (f). Letters next to each trend line represent the statistical relatedness of the FVs for each treatment: Student’s paired *t*.test; *P*<0.05; n=12.

In TuMV-GFP inoculated and systemic leaves, the FVs and CP levels were similar in *bag7* and Col-0 plants (Figure 3d, e and f; P<0.05) demonstrating that BAG7 does not restrict local or systemic TuMV-GFP infection.

### TGB3 and 6K2 induce *NAC089* and *NAC103* expression

The *NAC089* and *NAC103* encode transcription factors and are among the canonical UPR genes that relay ER stress signals from bZIP60 and bZIP28. NAC089 regulates downstream genes involved in programmed cell death and proteasome degradation of proteins, such as *BAG6,* while NAC103 activates downstream UPR genes including *CRT1* and *CNX1* and *PDI5* (Sun *et al.*, 2013, Yang *et al.*, 2014). To gain evidence that the downstream UPR-responsive genes *NAC089* and *NAC103* are sensitive to the potexvirus TGB3 or potyvirus 6K2 proteins, binary plasmids containing these PlAMV TGB3, PVX TGB3, TuMV 6K2 and PVY 6K2 genes were agro-infiltrated to Col-0 as well as *bzip60, bzip28* and *bzip17* plants. Then we performed qRT-PCR to examine changes in transcript accumulation at 2 dpi. Elevated levels of *NAC089* ranged between 8- and 37-fold in Col-0 plants treated with each viral elicitor (Figure 4a; P<0.05). However, in the *bzip60, bzip28*, and *bzip17* plants, the *NAC089* transcripts were reduced (Figure 4a; P<0.05). In *bzip60/bzip17* and *bzip60/bzip28* plants, there was no induction following treatment with each of these viral elicitors, suggesting an additive effect of the combined genes.

**Figure 4.**
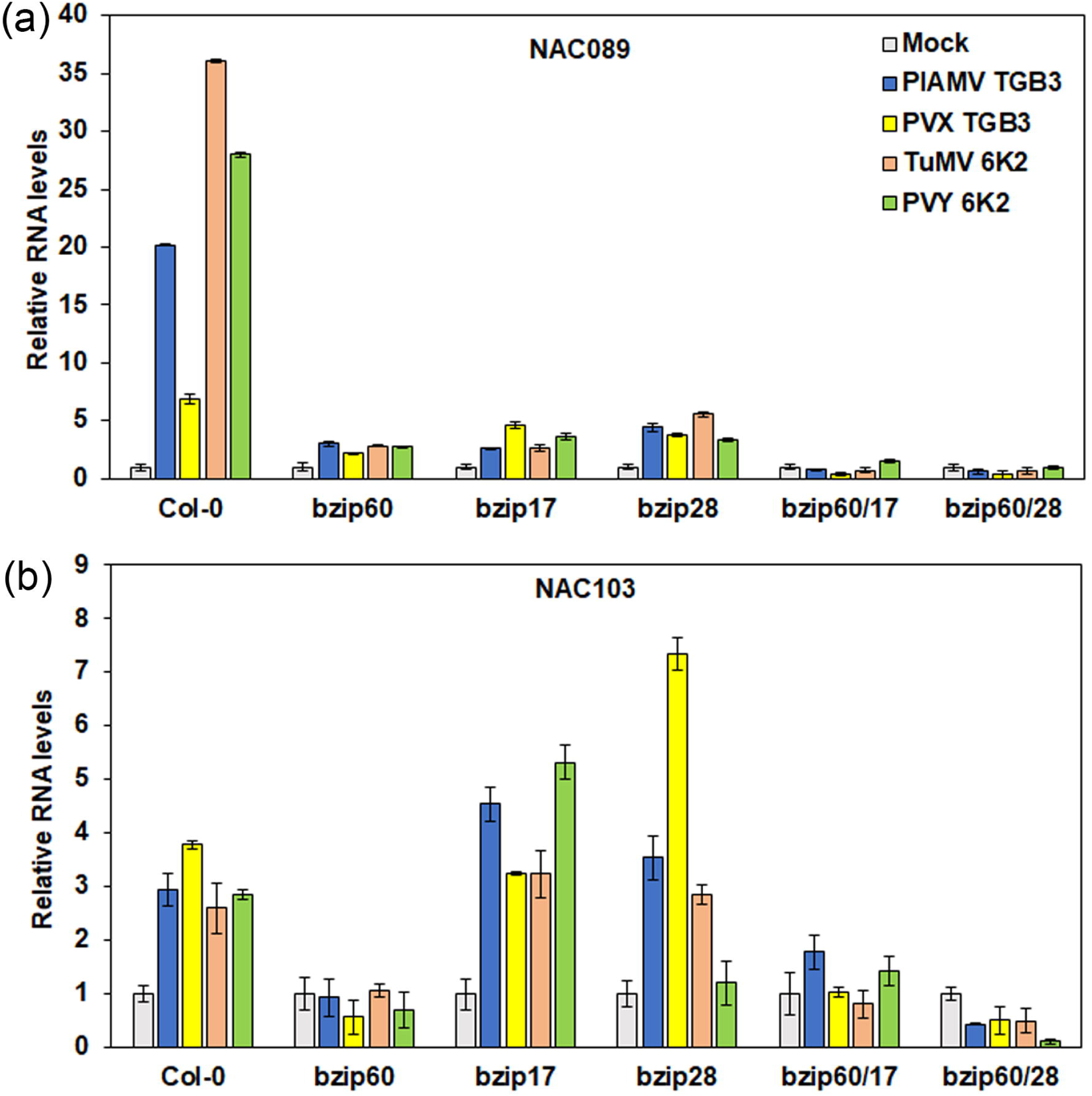
Overexpression of *NAC089* and *NAC103* in wild-type and bZIP mutants *Arabidopsis* leaves following agro-delivery of TGB3 and 6K2 genes. Bar graphs depict the average relative NAC089 (a) and NAC103 (b) transcript levels in samples harvested at 2 dpi. Agro-delivery of each viral factor is identified in the chart legend in panel (a). Error bars represent SD and asterisks indicate significant differences between treatments with viral factors and the mock: Student’s *t*.test; *P*<0.05; n=3.

In Col-0 plants, *NAC103* was induced by approximately 2.5-fold following expression of the TGB3 or 6K2 proteins (Figure 4b; P<0.05). In *bzip17* and *bzip28* plants, *NAC103* expression ranged from 3- to 8-fold above the control leaves, except in the case of PVY 6K2 which did not show a significant change in *bzip28* plants. Gene induction was not observed in *bzip60, bzip60/bzip17* and *bzip60/bzip28* plants (Figure 4b; P<0.05). These combined data suggest that TGB3 and 6K2 induced *NAC089* and *NAC103* expression.

### NAC089, but not BAG6, reduces the systemic accumulation of PlAMV-GFP

NAC089 relays UPR gene activation from bZIP60, bZIP28, and bZIP17, to upregulate ER stress-responsive genes, including *BAG6* (Yang *et al.*, 2014). BAG6 is a co-chaperone that is conserved across eukaryotes involved in protein quality control and proteasome elimination of abnormal proteins (Yamamoto *et al.*, 2017). The human BAG6 aids the folding of aggregation-prone proteins, polyubiquitinated proteins, and transmembrane proteins (Kawahara *et al.*, 2013). The Arabidopsis BAG6 contributes to autophagy that is associated with disease resistance to *Botrytis cinerea* (Li and Dickman, 2016, Li *et al.*, 2016). We observed the accumulation of PlAMV-GFP and TuMV-GFP in the systemic leaves of *nac089* and *bag6* plants and quantified FVs (Figure 5; n=6). We did not conduct experiments to test the role of *NAC103* in virus infection because there are no available homozygous mutant lines (https://abrc.osu.edu) with altered *NAC103* expression.

**Figure 5.**
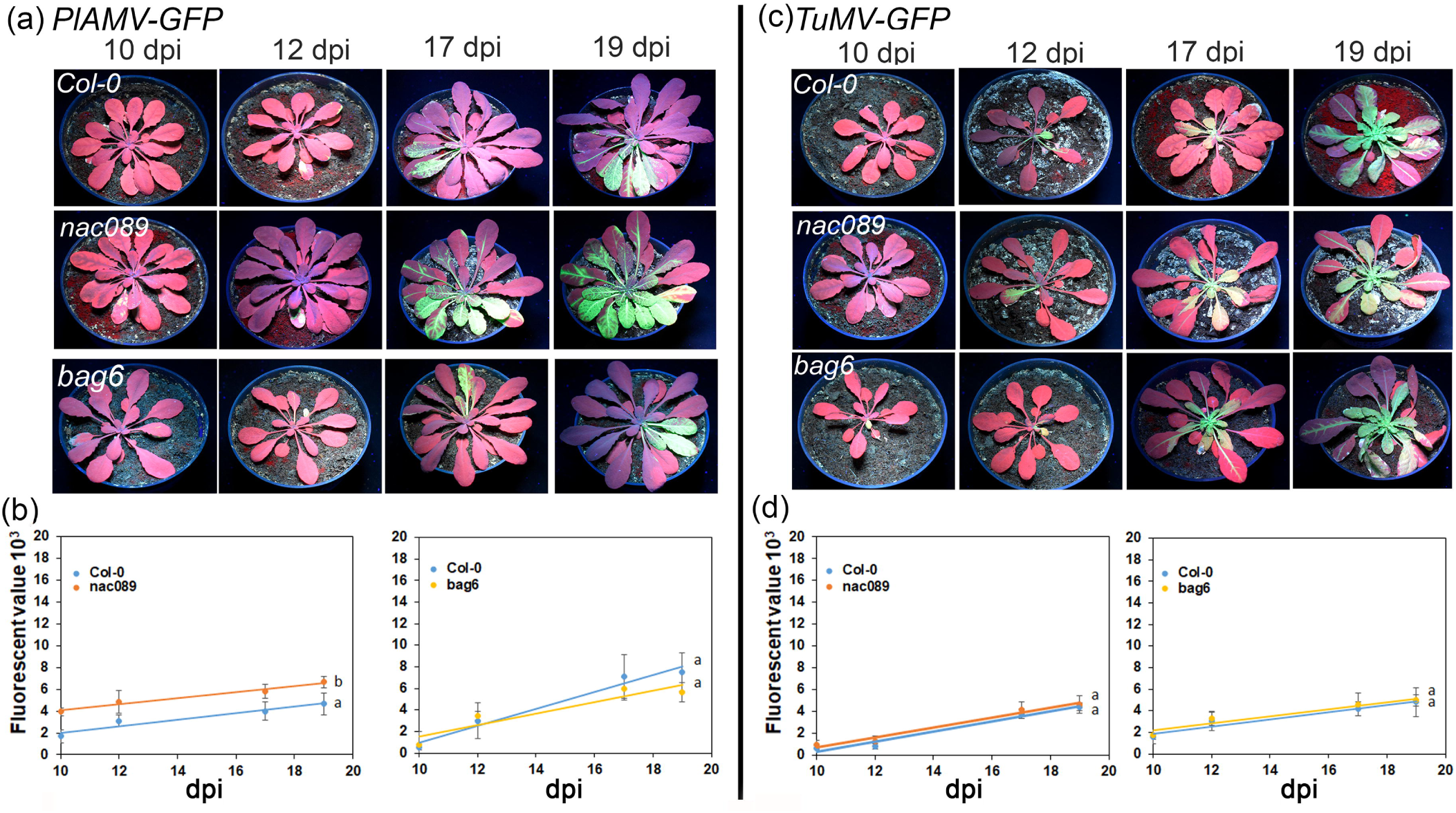
Local and systemic infection of PlAMV-GFP and TuMV-GFP in *nac089* and *bag6*, and Col-0 plants. Representative images of Col-0, *nac089*, and *bag6* plants infected with PlAMV-GFP (a) or TuMV-GFP (d). Scatter plot values and trend lines represent the average FVs and SDs for each mutant line and Col-0 inoculated with PlAMV-GFP (b) and TuMV-GFP (e). Letters next to each trend line represent the statistical relatedness of the FVs for each treatment: Student’s paired *t*.test; *P*<0.05; n=6. Percentage of plants that are infected by PlAMV-GFP (c) or TuMV-GFP (f) at 10 to 19 dpi.

As in previous experiments, GFP was a reporter of PlAMV-GFP and TuMV-GFP systemic infection in *nac089* and *bag6* plants. Among PlAMV-GFP inoculated plants, the FVs were significantly higher in *nac089* than in Col-0 plants (Figure 5b; P<0.05). However, the average FVs in Col-0 and *bag6* plants were not significantly different (Figure 5b; P>0.05). These results suggest that NAC089, like bZIP60 and bZIP17, although to a lesser extent, reduce PlAMV-GFP systemic accumulation, whereas BAG6 is not a factor of PlAMV-GFP infection. Regarding TuMV-GFP infection, the FVs in systemic leaves were unaffected in *nac089, bag6* compared to Col-0 plants (Figure 5d; P>0.05). Finally, these results suggest that NAC089 has a role in restricting PlAMV-GFP systemic accumulation and that BAG6 is not involved in the systemic accumulation of either virus.

### Increased protein folding capacity reduces PlAMV-GFP accumulation

Cells gain tolerance to ER stress by increasing the protein folding capacity of the ER through enhanced expression of chaperones. BiPs are among the primary targets of UPR signaling pathways converging on their activation in response to ER stress. Because *Arabidopsis BiP1* and *BiP2* share 97% nucleotide and 99% amino acid sequence identities, we could not sperately quantify these transcript levels using qRT-PCR. BiP3 is 80% identical to BiP1/2, and PCR primers can differentially detect these transcripts (Noh *et al.*, 2003, Srivastava *et al.*, 2013). At 5 dpi, the *BiP1/2* transcripts were higher following agro-delivery of TuMV or PVY *6K2* (Figure 6a; P<0.05) but not following expression of the PlAMV or PVX *TGB3* genes. However, *BiP3* transcripts were elevated 3- to 4-fold above the control in response to expression of PlAMV TGB3, PVX TGB3, PVY 6K2, and TuMV 6K2 (Figure 6a; P<0.05).

**Figure 6.**
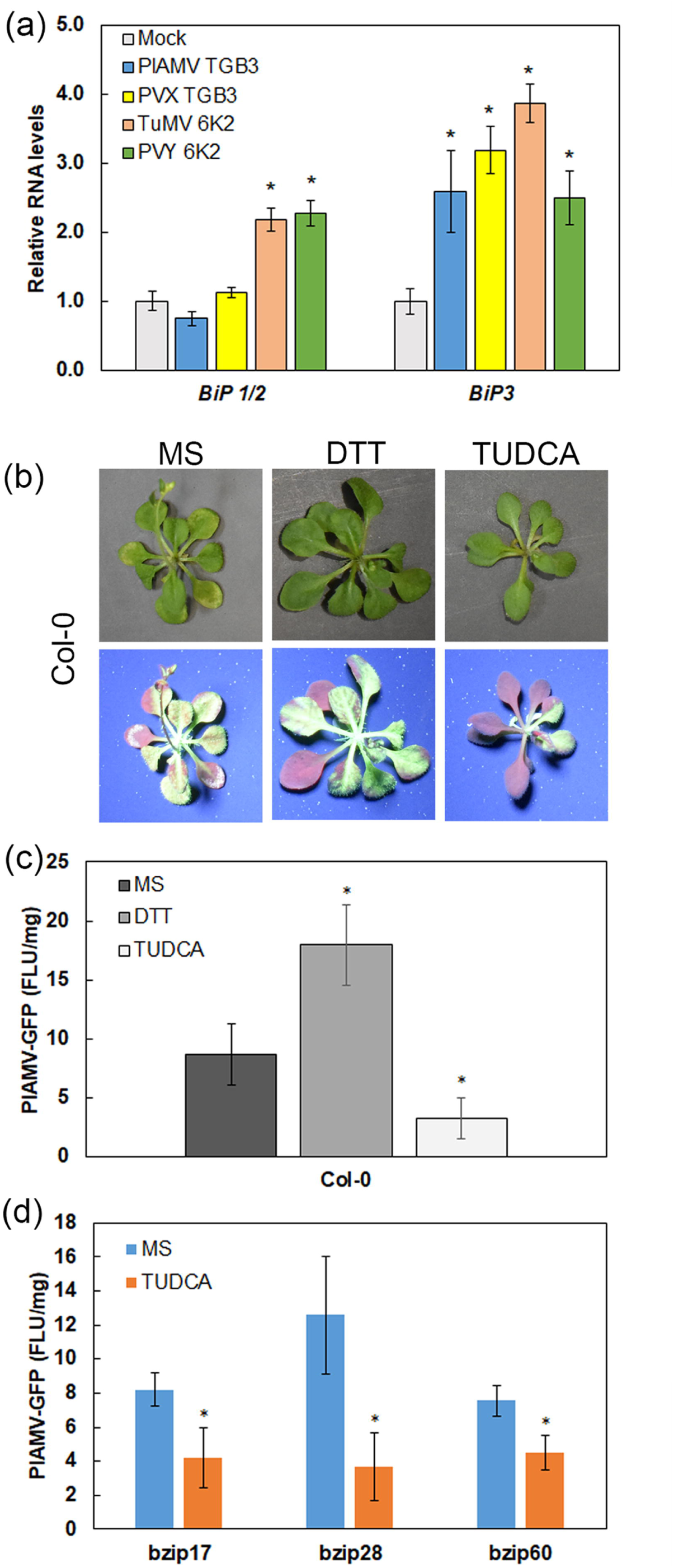
TUDCA represses PlAMV-GFP accumulation. qRT-PCR results are providing relative RNA levels of *BiP1/2* (a), *BiP3* (b) following treatments with each viral factor identified by color in the chart legend. Error bars represent SD and asterisks indicate significant differences to the mock control: Student’s *t*-test; *P*<0.05; (n=3). (b) Bright-field and fluorescent images of Col-0 plantlets grown on MS medium, 0.1mM of DTT or 0.5mM of TUDCA containing medium and infected with PlAMV-GFP at 15 dpi. (c) Fluorescence from PlAMV-GFP infected Col-0 plantlets grown on MS medium, 0.1mM of DTT or 0.5mM of TUDCA containing medium at 15 dpi. The results are expressed in arbitrary fluorescence units per fresh weight (FLU/mg). The error bars represent SD and asterisks indicate significant difference from plantlet on MS medium: Student’s *t*.test; *P*<0.05; n=3-4. (e) Fluorescence from PlAMV-GFP infected *bzip17, bzip28* and *bzip60* mutant plantlets grown on MS medium or 0.5mM of TUDCA containing medium at 15 dpi (FLU/mg). The error bars represent SD and asterisks indicate a significant difference between treatment and non-treated samples: Student’s *t*-test; *P*<0.05; n=3-4.

To examine the importance of the protein folding capacity in virus accumulation, we quantified virus-GFP fluorescence in infected plantlets grown on MS medium with added DTT and TUDCA. DTT causes significant ER stress in plant cells by reducing protein disulfide bond formation and reducing the protein folding capacity. TUDCA is a chemical chaperone that alleviates ER stress when applied to plants because it mitigates protein aggregation and stabilizes protein conformation (Zhang *et al.*, 2015, Fernández-Bautista *et al.*, 2017, Uppala *et al.*, 2017). We were unable to infect young plantlets with TuMV-GFP and conducted these experiments only with PlAMV-GFP. We inoculated 10-day-old Col-0 seedlings with PlAMV-GFP and transferred them to MS medium alone, with added 0.1 mM DTT, or with added 0.5 mM TUDCA (Figure 6b and c). At 15 dpi, we measured the fluorescence in crude extracts and calculated the average FV relative to the average sample fresh weight for each treatment (Figure 6c). The FV of PlAMV-GFP infected plantlets grown on DTT containing medium were higher than FV of infected plantlets grown on MS medium alone (Figure 6c; P<0.05). These data suggest that compromising the protein folding capacity of the cell favors virus infection. On the other hand, PlAMV infected plantlets grown on the TUDCA containing medium showed lower FV than the controls (Figure 6c; P<0.05) suggesting that increasing the protein folding capacity leads to a decrease in virus accumulation.

We also grew *bzip17, bzip28* and *bzip60* infected plantlets with PlAMV-GFP on MS media alone, with 0.1 mM DTT or with 0.5 mM TUDCA. These infected plantlets grew slowly on DTT containing medium and we could not generate an adequate fluorescence dataset. Thus, we only analyzed infected seedlings grown on MS and TUDCA containing medium. PlAMV-GFP fluorescence was lower in *bzip17, bzip28*, and *bzip60* plants grown on TUDCA medium compared to the fluorescence obtained from plants grown in regular MS media (Figure 6d; P<0.05). These combined data demonstrate that TUDCA can mitigate mutations affecting the UPR machinery during PlAMV-GFP infection. Finally, this result shows that expanding the protein folding capacity leads to decreased PlAMV-GFP accumulation and suggests that the maintenance of the functional protein folding machinery is vital for plants to restrict virus infection.

## Discussion

This study demonstrates that the bZIP17/bZIP28 pathway, alongside the IRE1/bZIP60 pathways, restrict PlAMV and TuMV infection in Arabidopsis plants. The PlAMV TGB3 and TuMV 6K2 proteins trigger the bZIP28/bZIP17 pathway and the IRE1/bZIP60 pathway of the UPR, which increases the expression of NAC089 transcription factors. These alternative pathways converge to activate expression of chaperones like BiPs, that are essential for the proper function of the cell, but also restrict virus infection.

Data presented here demonstrates bZIP60, bZIP17, and BAG7 serve to limit early events in PlAMV-GFP infection. The higher accumulation of PlAMV-GFP in *bzip17* and *bzip60* than in Col-0 plants, clearly demonstrated that both bZIP17 and bZIP60 have a role in restricting local and systemic infection. In Figure 2 PlAMV-GFP accumulation in the inoculated and systemic leaves in the *bzip17* and *bzip60* plants were not significantly different, indicating that both factors are essential for restrict PlAMV accumulation. Moreover, systemic infection was not statistically different in the *bzip60/bzip17* relative to the *bzip60* plants indicating that these factors do not act additively. Since bZIP60 and bZIP17 form heterodimers for inducing target genes (Deppmann *et al.*, 2004), these two factors may act synergistically to induce specific genes that restrict PlAMV-GFP accumulation. We were surprised by the results indicating that PlAMV-GFP accumulation is unaltered mainly in *bzip28* and *bzip28/bzip60* compared to Col-0 plants. Since PlAMV-GFP CP and FVs are typically higher in *bzip60* plants, we expected to observe higher PlAMV-GFP accumulation in both b*zip60/bzip28* and *bzip60* plants. It is possible that bZIP28 and bZIP60 redundantly upregulate unknown factor(s) providing positive support for PlAMV infection and that defects in both genes overcome the defect in *bzip60* plants. Such factors would likely not be downstream of NAC089 (which also serves to limit PlAMV infection) but would likely involve another downstream signal.

Given that it is well established that the bZIP17/bZIP28 and IRE1/bZIP60 pathways relay information for nuclear gene expression that activate pro-survival and pro-death pathways in plants, it is reasonable to consider that these ER stress sensors might also initiate signaling cascades that can activate cellular responses that may limit or promote virus infection. In this model, bZIP60 and bZIP17 synergistically induce genes restricting PlAMV infection, whereas bZIP60 and bZIP28 independently induce genes supporting PlAMV infection (Figure S2). In *bzip60* and *bzip17* plants, bZIP17 or bZIP60 alone do not induce genes restricting virus infection, whereas bZIP28 induces genes that promote higher PlAMV accumulation. In Col-0 *and bzip28* plants, bZIP60 and bZIP17 may together induce genes restricting virus accumulation, while bZIP60 may act alone to induce genes supporting PlAMV infection. Finally, in *bzip60/bzip28* plants, the phenotype observed in *bzip60* plants is alleviated by the absence of supporting genes induction by bZIP28 (Figure S2).

This study also showed that bZIP60 and bZIP28 are principally important for limiting TuMV-GFP accumulation in the inoculated and systemic leaves (Figure S3). The results in Figure 2 indicate that bZIP28 and bZIP60 are essential, although not additive, to limit TuMV-GFP accumulation. The GFP FVs measured in systemic leaves were comparable between *bzip60, bzip2*8, and *bzip60/bzip28* plants, suggesting that these factors may synergistically activate genes that limit infection. The bZIP17 and NAC089 likely do not contribute to the restriction of TuMV-GFP accumulation since GFP FVs were unaltered in the *bzip17 or nac089* plants compared to Col-0 plants. Based on these results we propose a model in which bZIP60 and bZIP28, but not bZIP17 are virus-limiting factors, that may have overlapping target genes (Figure S3).

The bZIP28 resides in a bZIP28/BAG7/BiPs complex in the ER under normal conditions, and this complex dissociates during ER stress, sending bZIP28 and BAG7 into the nucleus where they separately regulate gene expression (Li *et al.*, 2017). Surprisingly loss of *BAG7* seems to hamper PlAMV-GFP infection in the inoculated leaves whereas bZIP28 is not a factor during PlAMV infection. Further experiments are needed to test whether stress tolerance associated with BAG7 is the result of BAG7 chaperone functions in the ER as well as BAG7-WRKY29 interactions in the nucleus. The hypothesis that *BAG7-WRKY29* gene regulation is a factor in virus infection is particularly intriguing because WRKY29 is engaged in pattern-triggered immunity (Asai *et al.*, 2002). Overexpressing WRKY29 may enhance disease resistance to *Fusarium graminearum* infection and other pathogens (Sarowar *et al.*, 2019). It is reasonable to consider that the PlAMV TGB3 protein might activate UPR in a manner that produces cytoprotective chaperones while also managing anti-viral immunity. We also postulated that ER-to-nucleus signaling via the bZIP28 branch of the UPR would be less restricted in *bag7* plants (Williams *et al.*, 2010). Notably, loss of *BAG7* expression did not alter the levels of TuMV-GFP in systemic leaves suggesting the difference between bZIP17 and bZIP28 in potyvirus and potexvirus restriction is linked to their own activation. Finally, for both PlAMV-GFP and TuMV-GFP BAG7 seems to act independent of bZIP28 in regulating virus infection.

Other research shows that bZIP transcription factors form heterodimers to expand the diversity of ER stress-induced genes that they can activate. Many reports have shown that bZIP17, bZIP28, and bZIP60 co-regulate specific genes in a manner that can expand the pattern of tissue expression, the timing of gene induction, or the magnitude of the response (Liu and Howell, 2010, Henriquez-Valencia *et al.*, 2015, Angelos *et al.*, 2017, Kim *et al.*, 2018, Ruberti *et al.*, 2018). These bZIP transcription factors can also act alone, as homodimers, to activate specific downstream genes. For example, bZIP60 directly binds to the *cis*-element pUPRE-III (Sun *et al.*, 2013), whereas only bZIP17 can activate some specific downstream transcription responses in response to salt stress (Liu *et al.*, 2007b). Our results support a model in which bZIP60 act independent of bZIP28 for support PlAMV infection or, can act synergistically with bZIP17 or bZIP28 to limit PlAMV and TuMV infection, respectively (Figure 7). Consequently, the different affinities of bZIP17, bZIP28 and bZIP60 to *cis*-regulatory elements of UPR genes could explain the differences observed between PlAMV and TuMV infection in the KO mutant plants and reveals a fine-tuning of UPR in plants following virus infection.

**Figure 7.**
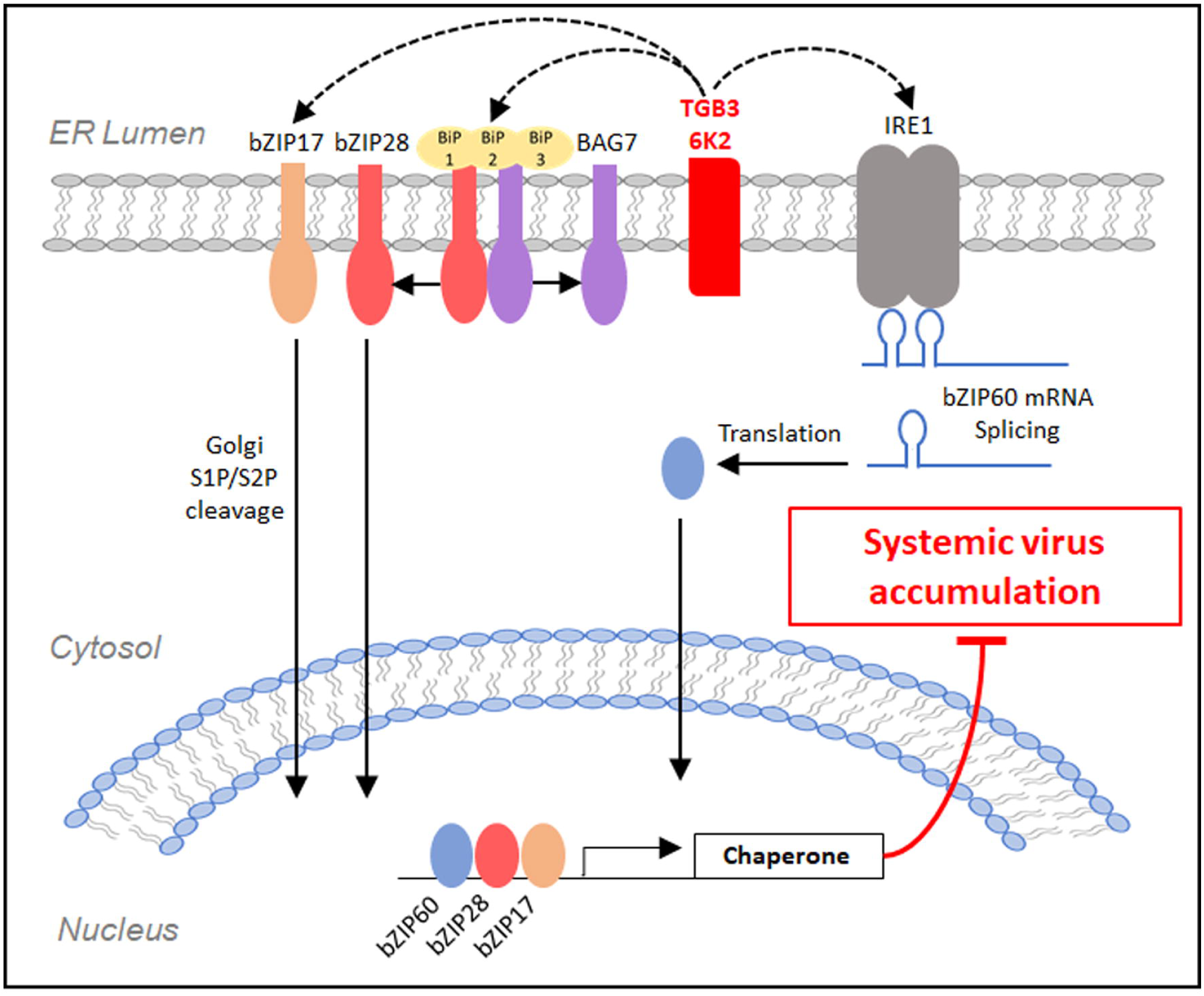
Proposed model for the function of UPR during virus infection. During poty- and potexvirus infection, the 6K2 and TGB3 viral proteins, respectively, induce the two arms of UPR. The IRE1 splices the *bZIP60* mRNA producing a truncated bZIP60 transcription factor. bZIP17 and bZIP28 move to the Golgi apparatus, S1P and S2P release the cytosolic part which goes to the nucleus. Once in the nucleus, bZIP17, bZIP28 and bZIP60 induce transcription of chaperone proteins that increase the ER folding capacity leading to repression of virus accumulation.

Most reported studies of UPR signaling in Arabidopsis have typically used ER stress-inducing agents such as heat, tunicamycin, or DTT. Investigations to uncover the molecular basis of adaptive UPR typically examine root and shoot growth of seedlings following short term or chronic treatments with these ER stress-inducing agents. The results of such studies indicate that bZIP28 and bZIP60 act in parallel to modulate common sets of ER stress-responsive genes engaged in adaptive UPR and ER stress recovery as well as chronic ER stress (Angelos *et al.*, 2017, Ruberti *et al.*, 2018). Prior studies characterized bZIP28 and bZIP60 as having partially redundant roles in UPR and stress recovery in roots, but function in parallel in shoot tissues (Ruberti *et al.*, 2018). By using identified viral elicitors of UPR and by infecting Arabidopsis plants with PlAMV-GFP and TuMV-GFP, we were able to contrast the transcriptional responses to ER stress and the requirements for bZIP factors to limit virus accumulation. Until now, bZIP17 was only known for its ability to engage in salt stress responses and vegetative growth. This study reports the involvement of bZIP17 in virus-induced ER stress and suggests that bZIP17 is also involved in maintaining ER homeostasis under chronic virus-induced ER stress (Figure 7).

This study also examined the roles of two NAC factors that are known to relay information from the bZIP factors for UPR gene activation (Nawkar *et al.*, 2018). The results indicate that *NAC089* and *NAC103* respond to TGB3 and 6K2 transient expression. PlAMV-GFP, but not TuMV-GFP accumulation was elevated in *nac089* than in Col-0 plants, suggesting that NAC089 restricts PlAMV infection. NAC089 modulates *BAG6* gene expression, and our data shows that BAG6 is not a factor in regulating PlAMV-GFP and TuMV-GFP accumulation. BAG6 was particularly interesting because of its function as a co-chaperone engaged in protein turnover in mammals and its role in plants in regulating autophagy and programmed cell death in response to fungal pathogens (Yang *et al.*, 2014, Li and Dickman, 2016, Li *et al.*, 2016). We carried out a brief analysis of ATG8 lipidation in *ire1a/ire1b, bzip60, bzip28, bzip17, nac089* and *bag6* plants which did not reveal differences in autophagy processes following PlAMV-GFP and TuMV-GFP infection suggesting that autophagy is not linked to UPR during these infection (Figure. S4). Overall, the results in this study suggest that the role of NAC089 during PlAMV infection is probably linked to regulating ER stress and cellular homeostasis rather than autophagy and cell death.

The UPR is mainly responsible for increasing the protein folding capacity of the ER through enhanced expression of the chaperones such as *BiP, CRT, CNX*, and *PDI*. We previously demonstrated that TGB3 activates expression of *CRT2* and *PDI* in Arabidopsis and *N. benthamiana* (Ye *et al.*, 2011) and this study showed that TGB3 and 6K2 induce expression of *BiPs*. Overexpression of *N. tabacum BiP-like protein 4* protects against viral-induced necrosis (Leborgne-Castel *et al.*, 1999, Ye and Verchot, 2011, Ye *et al.*, 2013). We show that decreasing or increasing the folding capacity of Arabidopsis plants through DTT and TUDCA application leads to a significant increase or decrease of PlAMV-GFP accumulation, respectively. Using TUDCA to mitigate defects in UPR signaling during PlAMV infection provided further evidence that maintaining healthy protein folding activities is essential for limiting virus infection. Altogether, these results suggest that during virus infection, UPR works to maintain a functional protein folding machinery for repress virus accumulation (Figure 7). The role of the protein folding machinery in virus inhibition is not clear, but one can imagine that the virus actively highjacks the protein folding machinery from its normal focus on cellular proteins to fold viral proteins. Under these conditions, increasing the protein folding capacity would allow the cell to protect homeostatic protein production and ensure proper production of defense proteins. However, regarding the synergic action of bZIP factors involve in UPR associated virus inhibition and differences observed between PlAMV and TuMV, the role of UPR is probably not limited to protein folding capacity and further experiments are needed to delineate the role of UPR in repressing virus accumulation.

In conclusion, this paper brings new knowledge on UPR and details the function of the Arabidopsis UPR signaling during potyvirus and potexvirus infection that could be useful for increase plant virus resistance. We provide evidence of the role of bZIP17 under chronic ER stress led by virus infection. We reveal that the two UPR arms seem to be mainly associated with virus inhibition, at the exception of downstream factors of bZIP28 during PlAMV infection and probably repress virus accumulation by increasing protein folding capacity.

### Experimental procedures

#### Plant materials

We obtained from the ABRC (The Ohio State University, Columbus, OH, USA) *Arabidopsis thaliana* ecotype Columbia-0 (Col-0) and knockout independent homozygous transfer DNA (T-DNA) insertion lines; *bzip17* (SALK_104326), *bzip28-2* (SALK_132285), bzip60-2 (SAIL_283_B03),*nac089* (SALK_201394), *bag6-1* (SALK_047959), *bag7* (Salk_065883), *bzip60-2/28-2, bzip60-2/17*, and *ire1a-2/1b-4.* We verified all lines by PCR and maintained them in the laboratory (Williams *et al.*, 2010, Gaguancela *et al.*, 2016, Li *et al.*, 2016). Arabidopsis plants were grown in a long day (16h) or short day (12h) photoperiod at 23°C.

#### Agrobacterium delivery of virus genes and infectious clones

We delivered Agrobacterium (GV3101) cultures harboring the pMDC32 and pGWB505 binary vectors containing PVY 6K2, TuMV 6K2, PVX TGB3 or PlAMV TGB3 genes by infiltration to the underside of Arabidopsis leaves as previously reported (Gaguancela *et al.*, 2016) (ThermoFisher, Richardson, TX, USA). The pGWB505 constructs have GFP fused to the 3’ end of each viral gene. Control Agrobacterium (referred to as mock) harboring the empty pMDC32 or the GFP expressing pXF7FNF2.0 were infiltrated to Arabidopsis leaves. The GFP-tagged infectious clones of TuMV and PlAMV were reported previously (Gaguancela *et al.*, 2016). We verified each viral cDNA insertion in each plasmid by sequencing. Culture suspensions harboring the relevant expression constructs were prepared using a solution of 10 mM MES-KOH (pH 5.6), 10 mM MgCl_2_, and 200 μM acetosyringone and adjusted to OD600 = 0.7 to 1.0. A 1-ml needle-free syringe was used to infiltrate the Agrobacterium suspensions into leaves of 3-week-old Arabidopsis seedlings grown under short-day conditions. Fig. S1 shows representative timelines for PlAMV-GFP and TuMV-GFP infection in Arabidopsis plants.

#### RNA extraction and real-time quantitative RT-PCR

We plunged leaf punches into liquid nitrogen and then extracted the RNA using the Maxwell LEV simplyRNA purification kit (Promega Corp., Madison WI, USA). The cDNA synthesis was carried out using the High-capacity cDNA reverse transcription kit (ThermoFisher, Richardson, TX, USA) and random primers. For the qPCR assays, we verified the efficiencies of all primers by endpoint PCR and gel electrophoresis. The transcript abundance was quantified using the Power SYBR Green II PCR master mix (ThermoFisher) in the ABI 7500 PCR and Step-One Plus or the QuantSudio 3 machines (Applied Biosystems, Foster City, CA, USA). The relative increase of cellular transcripts was calculated using the comparative cycle threshold (CT) method, which employs the equation 2^-ddCT^ (Livak and Schmittgen, 2001). The endogenous control was *ubiquitin-10 or 18S*. Statistical analysis was carried out using the Student t-test to validate the results.

#### Immunoblot analysis

For detecting viral proteins, immunoblot analysis was carried out using virus-infected inoculated leaves, harvested at 5 or 8 dpi for PlAMV-GFP or TuMV-GFP, respectively. PlAMV-GFP fluorescence appeared earlier than TuMV-GFP fluorescence in inoculated leaves (Fig. S1). Total protein was extracted using standard methods (Ye *et al.*, 2011) and quantified using the Pierce™ Coomassie Plus assay (ThermoFisher, Rockford, IL, USA). Then nine µg of protein per sample were loaded onto 15% SDS-PAGE gels. For ATG8 detection, we harvested systemically infected leaves at 25 dpi, and total protein was extracted using a buffer comprised of 50 mM NaH2PO4 (pH 7.0), 200 mM NaCl, 10 mM MgCl2, 10% glycerol, 0.2% β-mercaptoethanol and protease inhibitor cocktail (Sigma-Aldrich Co., St. Louis, MO, USA). Then for each sample, 15 µg of protein were loaded onto 15% SDS-PAGE gels containing 6M urea. In both experiments, the Trans-Blot® Turbo™ Transfer System (Bio-Rad Laboratories, Hercules, CA, USA) was used for electroblot transfer of proteins to PVDF membranes where were later probed with antisera detecting the viral CPs (Agdia, Elckhardt, IN, USA) or Atg8 (Abcam, Cambridge, UK).

#### Analysis of local and systemic infection

Analysis of local and systemic infection was optimized and reported (Gaguancela *et al.*, 2016). To analyze PlAMV-GFP or TuMV-GFP in the inoculated leaves, we acquired fluorescent images at 5 dpi and 8 dpi, respectively. To analyze systemic PlAMV-GFP or TuMV-GFPinfection, we acquired fluorescent images at intervals between 10 and 19 dpi. Image J software quantified the fluorescent intensity. The results represent the average fluorescence value (FV) for at least 6 plants. For comparisons of inoculated leaves, reported the average FV for each treatment relative to the control. We plotted the FV values representing systemic infection and used ANOVA (P<0.05) to validate the results.

#### Chaperone protection assays

Arabidopsis seeds were sterilized with 50% bleach and 1% Triton X-100 for 10 min and then washed five times with sterile water. We stratified seeds for three days at 4°C, and then germinated on solid (0.8% agar) ½-strength Murashige and Skoog medium including 2% glucose (½-MS) in a growth chamber for 10 days under long-day conditions. Seedlings were then vacuum infiltrated with the infectious clones of PlAMV-GFP (OD600=0.5 to 0.7) and transferred to liquid ½-MS medium for 24h. Seedlings were washed with ½-MS medium containing 100 µg/mL Timentin and then cultivated on solid ½-MS medium containing 100 µg/mL timentin plus 01.mM DTT or 0.5mM TUDCA. Plantlets were harvested at 15 dpi and ground in liquid nitrogen before GFP extraction in PBS buffer. We quantified GFP fluorescence using a Fluoroskan FL microplate fluorometer (Ex/Em=485 nm/538 nm), and the results were expressed in arbitrary fluorescence units per fresh weight (FLU/mg) and were averaged for three to four plants. Statistical analysis was conducted using the Student t-test (P<0.05).

## Acknowledgments

We thank Steven Howell for providing the homozygous mutant lines used in this study. This research was funded by NSF/USDA-AFRI joint award number 1759034.

## Supporting information

**Figure S1.** Representative images are capturing the timeline progression of PlAMV-GFP and TuMV-GFP infection in Col-0 plants.

**Figure S2.** Proposed model for PlAMV infection UPR in mutant plants.

**Figure S3.** Proposed model for TuMV infection UPR in mutant plants.

**Figure S4.** ATG8 lipidation following PlAMV-GFP and TuMV-GFP infection.

**Table S1.** Primers used in this study.

