## Supporting Information for "Multiple ER-to-nucleus stress signaling pathways become active during *Plantago asiatica mosaic virus* and *Turnip mosaic virus* infection in *Arabidopsis thaliana*"

**Supporting information detailed legends submitted as a separate Word document.**

**Figure S1.** Time course of PlAMV-GFP and TuMV-GFP infection in Col-0 plants.

Fluorescent images were acquired from 0 dpi to 19 after inoculation of PlAMV-GFP and TuMV-GFP in Col-0 plants. After the PlAMV-GFP inoculation, the GFP fluorescence appear at 4 dpi in infiltrated leaves (local infection; green boxes) and GFP fluorescence appear at 12 dpi in upper leaves (systemic infection; red boxes). After the TuMV-GFP inoculation, the GFP fluorescence weakly appear at 6 dpi in infiltrated leaves (local infection; green boxes) and GFP fluorescence appear at 12 dpi in upper leaves (systemic infection; red boxes).

**Figure S1.** Proposed model for PlAMV infection UPR in mutant plants.

In this model, bZIP28 and bZIP60 redundantly upregulate certain unknown factor(s) providing positive support for PlAMV infection (red box) and bZIP17 and bZIP60 synergistically upregulate factor restricting PlAMV infection (green box). In Col-o plants, bZIP60 and bZIP17 could synergistically induce genes restricting PlAMV infection whereas bZIP60 and bZIP28 could independently induce genes supporting PlAMV infection resulting in regular infection. In *bzip60* mutant plants, genes restricting virus are not well induced by bZIP17 alone (grey arrow) whereas genes supporting PlAMV infection are induce by bZIP28, resulting in high virus accumulation. In *bzip28* mutant plants, genes restricting and supporting PlAMV infection are both induced by bZIP60 resulting in regular virus accumulation. In *bzip60/bzip28* double mutant plants, the phenotype observed in *bzip60* plants is alleviated by the absence of supporting genes induction by bZIP28. In *bzip17* and *bzip60/bzip17* mutant plants, genes restricting virus are not induced whereas genes supporting PlAMV infection are induce by bZIP28, resulting in high virus accumulation.

**Figure S2.** Proposed model for TuMV infection UPR in mutant plants.

In this model, bZIP28 and bZIP60 synergistically upregulate factor restricting TuMV infection (green box) whereas bZIP17 is not a factor involve in TuMV infection. In Col-o plants, bZIP60 and bZIP28 could synergistically induce genes restricting TuMV infection resulting in regular infection. In *bzip60* and *bzip60/bzip17* mutant plants, genes restricting virus are not well induced by bZIP28 alone (grey arrow) resulting in high virus accumulation. In *bzip28* and *bzip60/bzip28* mutant plants, genes restricting virus are not induced resulting in high virus accumulation. In *bzip17* mutant plants, bZIP60 and bZIP28 could synergistically induce genes restricting TuMV infection resulting in regular infection.

**Figure S3.** ATG8 lipidation following PlAMV-GFP and TuMV-GFP infection.

Representative immunoblots detecting ATG8 and the phosphatidylethanolamine bound ATG8 (ATG8-PE; identified by arrowheads on the right) from systemic leave of Col-0, *ire1a/ire1b, bzip60, bzip28, bzip17, nac089* and *bag6* plants infected by PlAMV-GFP and TuMV-GFP. GFP blots confirm equal levels of virus infection and Coomassie blue stain (CBS) confirms equal protein loading among the lanes.
