## Supplementary figures and images for "Multiple ER-to-nucleus stress signaling pathways become active during *Plantago asiatica mosaic virus* and *Turnip mosaic virus* infection in *Arabidopsis thaliana*"

### Supplemental Figure 1

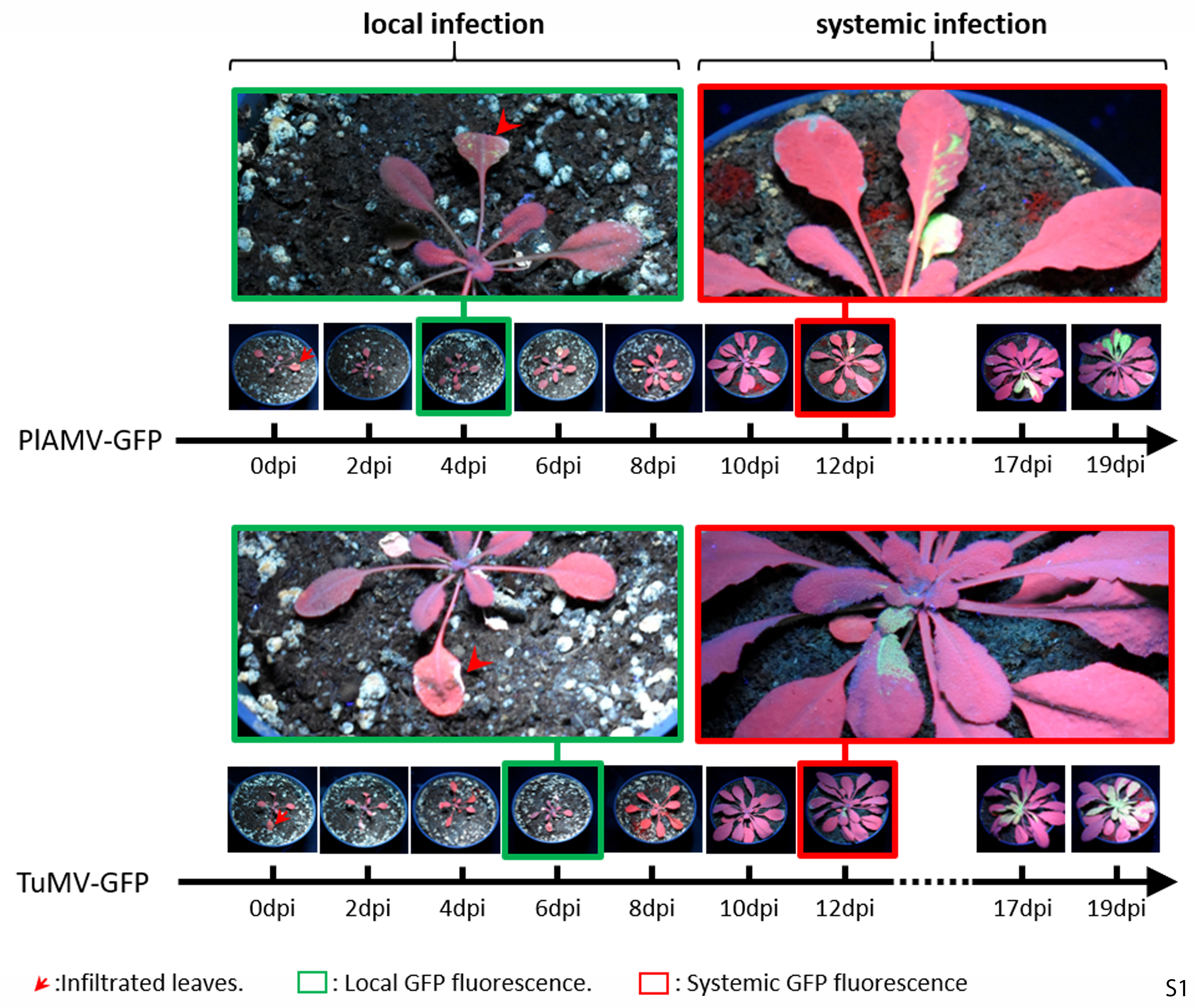

### Supplemental Figure 2

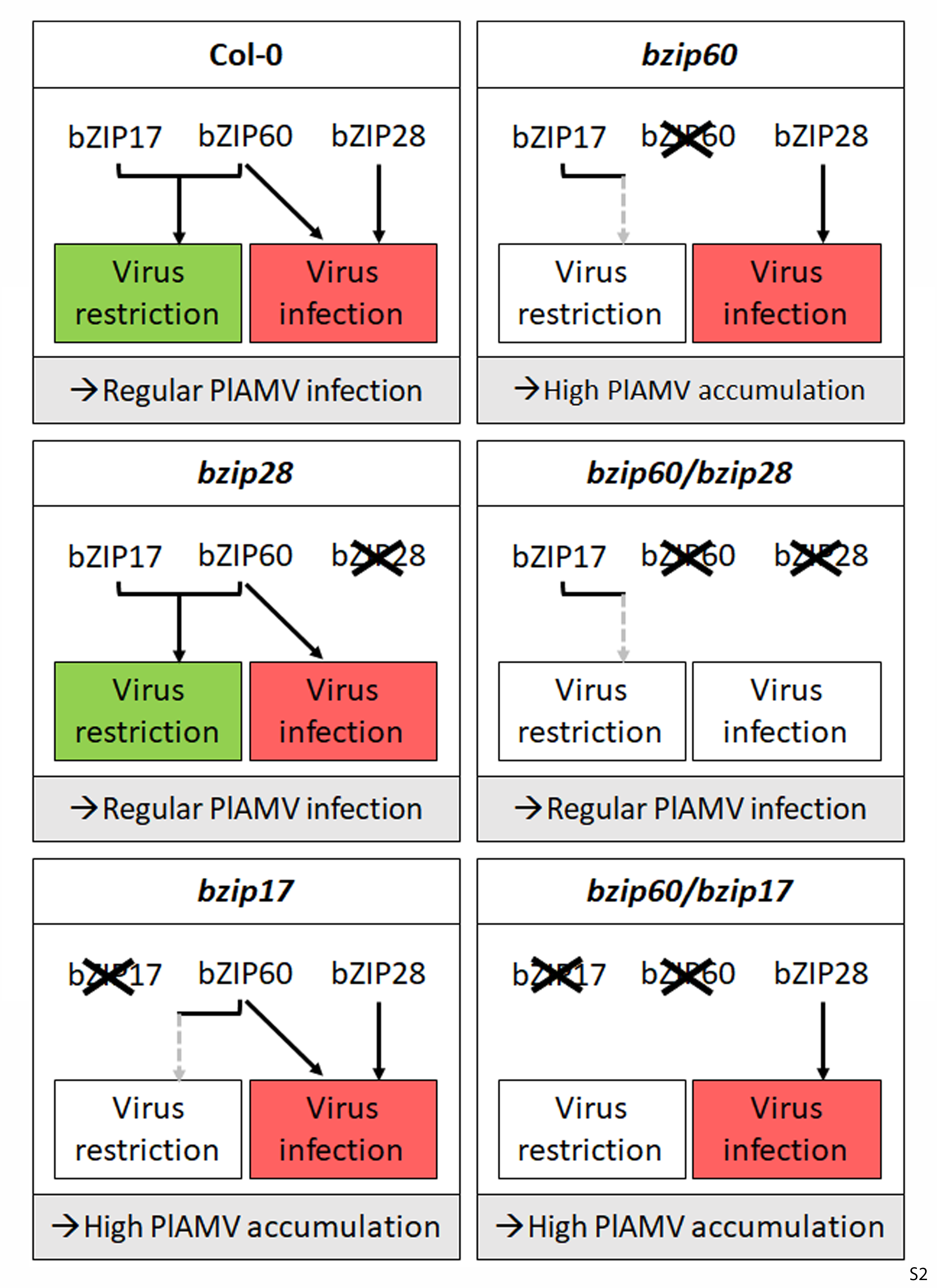

### Supplemental Figure 3

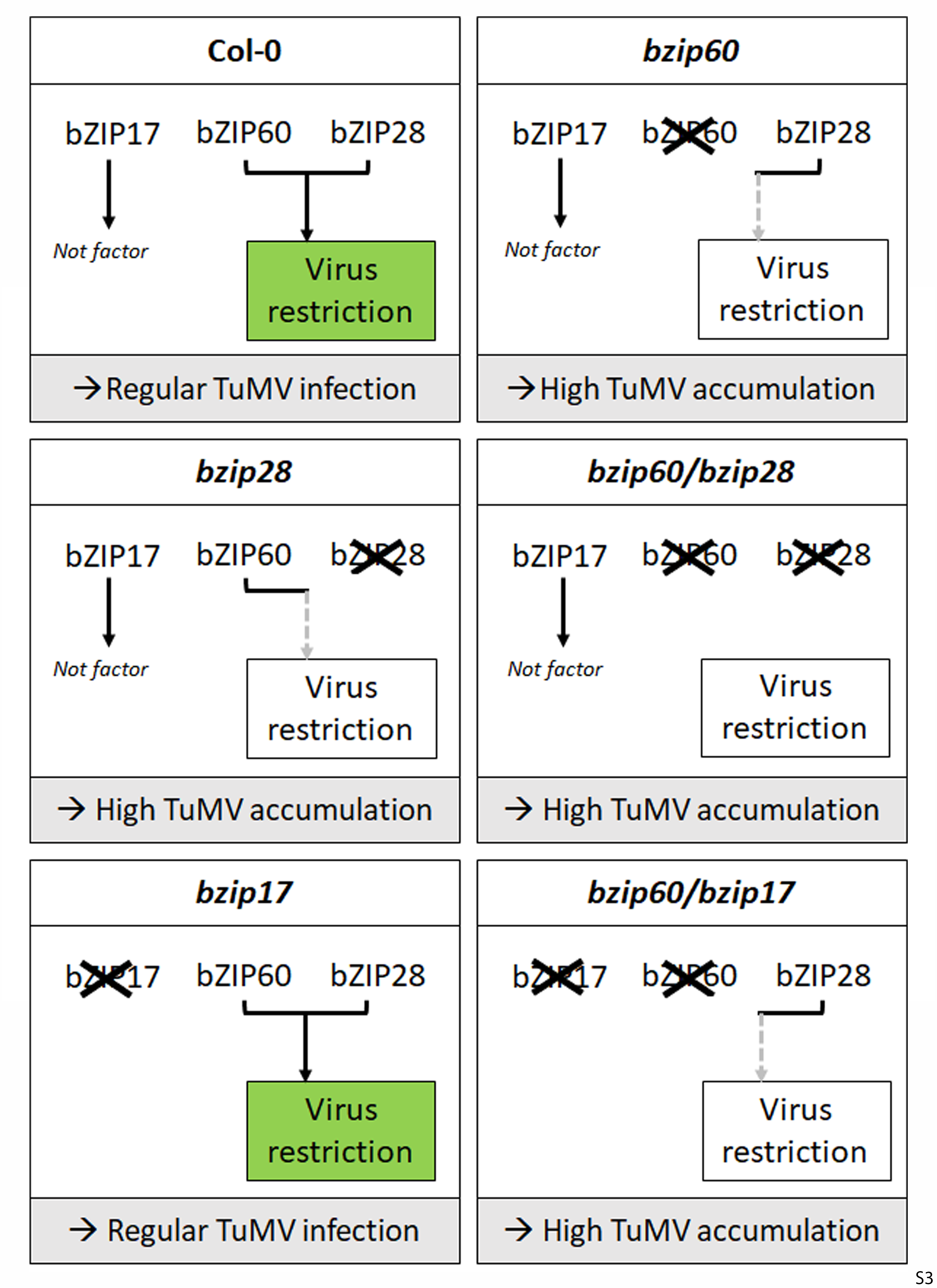

### Supplemental Figure 4

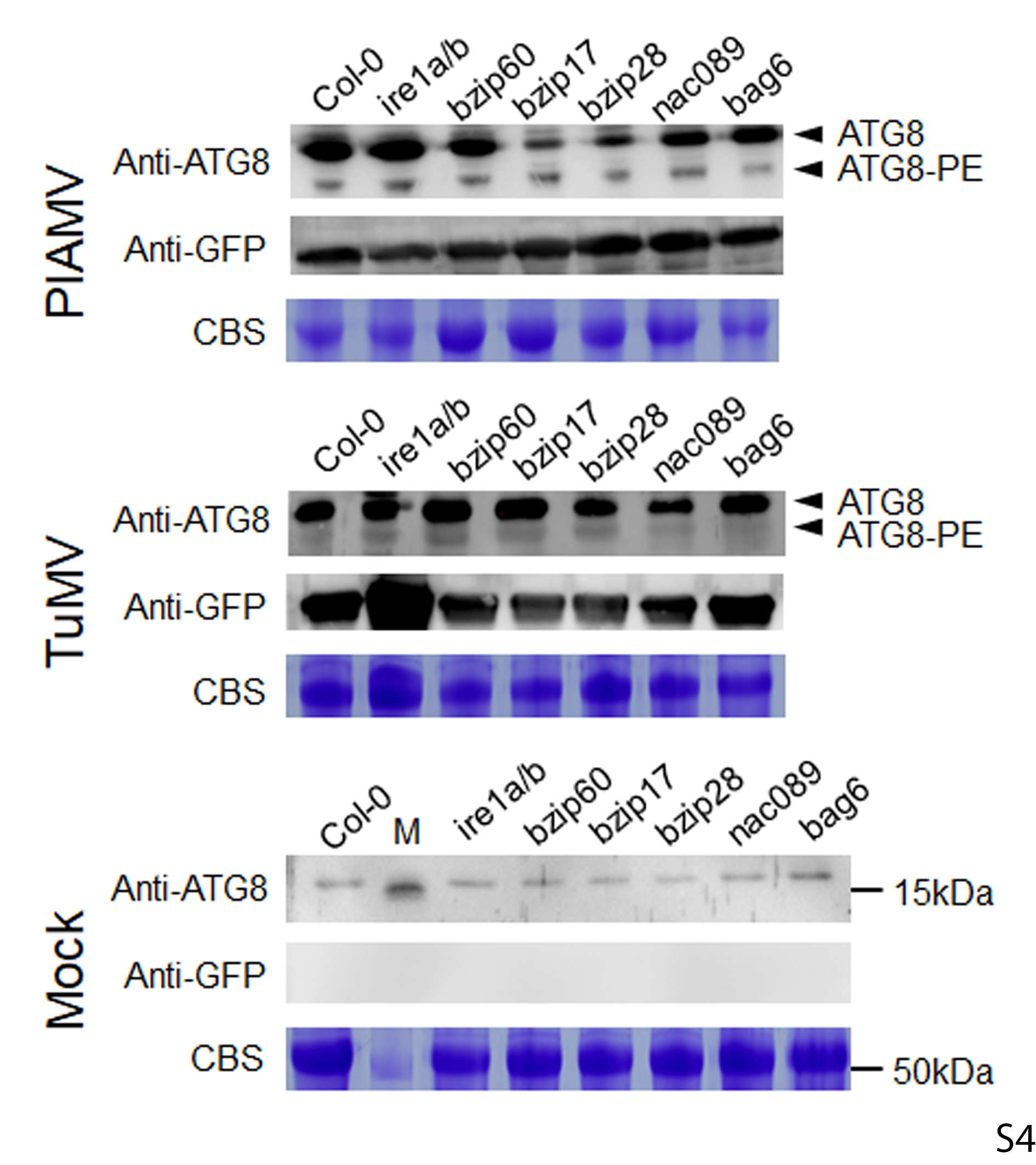
