## Supplemental Table 1 for "Multiple ER-to-nucleus stress signaling pathways become active during *Plantago asiatica mosaic virus* and *Turnip mosaic virus* infection in *Arabidopsis thaliana*"

| Table S1 |  |  |
| --- | --- | --- |
| **Primer name** | **Primer sequence 5' - 3'** | **Description** |
| AtbZIP17 FWD Set3 | CCTATGGCTCCAATGCCTTAT | NM_129659.3 Arabidopsis thaliana Basic-leucine zipper (bZIP) transcription factor family protein (BZIP17), mRNA |
| AtbZIP17 REV Set3 | GGATGTTCCAAGGGTGTTCT |  |
| AtbZIP60 F | CGCTGATGATTCCGGGAAGGAGAA | NM_103458.3 Arabidopsis thaliana basic region/leucine zipper motif 60 (BZIP60), mRNA |
| AtbZIP60 R | CTTCCTCTCTCTCGATCTAACCGC |  |
| AtbZIP28 F | GGCAGATTAGGAACCGTGAAA | NM_111917.5 Arabidopsis thaliana Basic-leucine zipper (bZIP) transcription factor family protein (BZIP28), mRNA |
| AtbZIP28 R | AGCGACATTCTCAGCCATAAC |  |
| AtNAC103 F | CAAAGGCGGTAAGGCAAACC | NM_125802.3 Arabidopsis thaliana NAC domain containing protein 103 (NAC103), mRNA |
| AtNAC103 R | CCGCGAGGTATTTTCCCGTA |  |
| AtNAC089 F | CTGATAGCGAGTGGTTCTTCTT | NM_122134.4 Arabidopsis thaliana NAC domain containing protein 89 (NAC089), mRNA |
| AtNAC089 R | TCCCAGTTGCTTTCCAGTATC |  |
| qPCR_AtBiP1/2 F | TCACTTGGGAGGTGAGGACTTT | NM_122737.4 Arabidopsis thaliana heat shock protein 70 (Hsp 70) family protein (BIP1), mRNA |
| qPCR_AtBiP1/2 R | CTCACATTCCCTTCGGAGCTTA | NM_180788.3 Arabidopsis thaliana Heat shock protein 70 (Hsp 70) family protein (BIP2), mRNA |
| qPCR_AtBIP3-F | CACGGTTCCAGCGTATTTCAAT | NM_001198015.2 Arabidopsis thaliana Heat shock protein 70 (Hsp 70) family protein (BIP3), mRNA |
| qPCR_AtBIP3-R | ATAAGCTATGGCAGCACCCGTT |  |
| AtUBQ10-F | GGCCTTGTATAATCCCTGATGAATAAG | NM_116771.5 Arabidopsis thaliana polyubiquitin 10 (UBQ10), mRNA |
| AtUBQ10-R | AAAGAGATAACAGGAACGGAAACATA |  |
| 18S F | ATGGCCGTTCTTAGTTGGTG | NR_141642.1 Arabidopsis thaliana 18S ribosomal RNA rRNA |
| 18S R | GTTAGCAGGCTGAGGTCTCG |  |
| PVX TGB3F | GGGGACAAGTTTGACGCAATCATACT | Used for RT-PCR verify gene expression following Agro-delivery |
| PVX TGB3R | GGGGACCACTTTGCCGTTCAAGGAG | Used for RT-PCR verify gene expression following Agro-delivery |
| PlAMV TGB3F | GGGGACAAGTTTGCTCCCACACGGAG | Used for RT-PCR verify gene expression following Agro-delivery |
| PlAMV TGB3R | GGGGACCACTTTGGAGCTTGGTTGAG | Used for RT-PCR verify gene expression following Agro-delivery |
| PVY 6K2F | GGGGACAAGTTTGAACAACTTCAC | Used for RT-PCR verify gene expression following Agro-delivery |
| PVY 6K2R | GGGGACCACTTTGCTGATTGAGTAAACC | Used for RT-PCR verify gene expression following Agro-delivery |
| TuMV 6K2F | GGGGACAAGTTTGAGCGACATGAGC | Used for RT-PCR verify gene expression following Agro-delivery |
| TuMV 6K2R | GGGGACCACTTTGGGTTACGGGTTCGG | Used for RT-PCR verify gene expression following Agro-delivery |
